# MeCP2 isoform e1 mutant mice recapitulate motor and metabolic phenotypes of Rett syndrome

**DOI:** 10.1101/357707

**Authors:** Janine M. LaSalle

**Affiliations:** UC Davis School of Medicine Department of Medical Microbiology and Immunology; UC Davis Genome Center; UC Davis MIND Institute; UC Davis School of Medicine Department of Psychiatry and Behavioral Sciences; UC Davis School of Medicine, Department of Public Health Sciences; UC Davis School of Veterinary Medicine, Department of Molecular Biosciences

## Abstract

Mutations in the X-linked gene *MECP2* cause the majority of Rett syndrome (RTT) cases. Two differentially spliced isoforms of exons 1 and 2 (MeCP2-e1 and MeCP2-e2) contribute to the diverse functions of MeCP2, but only mutations in exon 1, not exon 2, are observed in RTT. We previously described an isoform-specific MeCP2-e1 deficient male mouse model of a human RTT mutation that lacks MeCP2-e1 while preserving expression of MeCP2-e2. However, RTT patients are heterozygous females that exhibit delayed and progressive symptom onset beginning in late infancy, including neurologic as well as metabolic, immune, respiratory, and gastrointestinal phenotypes. Consequently, we conducted a longitudinal assessment of symptom development in MeCP2-e1 mutant females and males. A delayed and progressive onset of motor impairments was observed in both female and male MeCP2-e1 mutant mice, including hind limb clasping and motor deficits in gait and balance. Because these motor impairments were significantly impacted by age-dependent increases in body weight, we also investigated metabolic phenotypes at an early stage of disease progression. Both male and female MeCP2-e1 mutants exhibited significantly increased body fat compared to sex-matched wild-type littermates prior to weight differences. *Mecp2e1^-/y^* males exhibited significant metabolic phenotypes of hypoactivity, decreased energy expenditure, increased respiratory exchange ratio (RER), but decreased food intake compared to wildtype. Untargeted analysis of lipid metabolites demonstrated a distinguishable profile in MeCP2-e1 female mutant liver characterized by increased triglycerides. Together these results demonstrate that MeCP2-e1 mutation in mice of both sexes recapitulate early and progressive metabolic and motor phenotypes of human RTT.

## Introduction

Mutations in the X-linked gene encoding methyl CpG-binding protein 2 (MeCP2) cause the majority of Rett syndrome (RTT) cases. The specific role for MeCP2 in the molecular pathogenesis of RTT is complex, involving multiple molecular mechanisms and cell types. MeCP2 has two alternatively spliced forms (MeCP2-e1 and MeCP2-e2), and mutations in *MECP2e1* (exon 1) but not *MECP2e2* (exon 2) have been linked to RTT. The two isoforms differ in their N-terminals by alterative inclusion of *Mecp2 exon 2* amino acids. MeCP2-e1 contains amino acids from exons 1, 3, and 4 while MeCP2-e2 contains amino acids encoded by all four exons but utilizes an alternative start codon in exon 2, resulting in the translation of only exons 2, 3 and 4 (1) (Figure 1A). The vast majority of RTT mutations occur in *MECP2* exons 3 and 4; however, mutations in exon 1, but not exon 2, have been identified (1–3). MeCP2-e1 is the primary isoform expressed in the central nervous system (CNS) (4) and has higher protein stability than MeCP2-e2 (5), further suggesting that MeCP2-e1 is the dominant isoform in brain. We recently demonstrated a critical role for MeCP2-e1 in the pathogenesis of RTT (5) by creating a novel mouse model based on an orthologous MeCP2-e1 translation start site mutation identified in patients with classic RTT (2, 3) (*Mecp2e1^-/y^* mice) (Figure 1A). This isoform specific model lacks MeCP2-e1 translation while preserving expression of the MeCP2-e2 isoform, allowing for the examination of the specific function of MeCP2-e1. Male *Mecp2e1^-/y^* mice recapitulate many RTT-like neurologic deficits (5) including development of hind limb clasping, motor deficits, and early lethality (5). These findings are in contrast to a mouse model with selective deletion of exon 2 that did not show neurologic symptoms (6). Together the evidence suggests that MeCP2-e1 deficiency alone is sufficient to induce neurologic symptoms underlying RTT in male mice.

**Figure 1.**
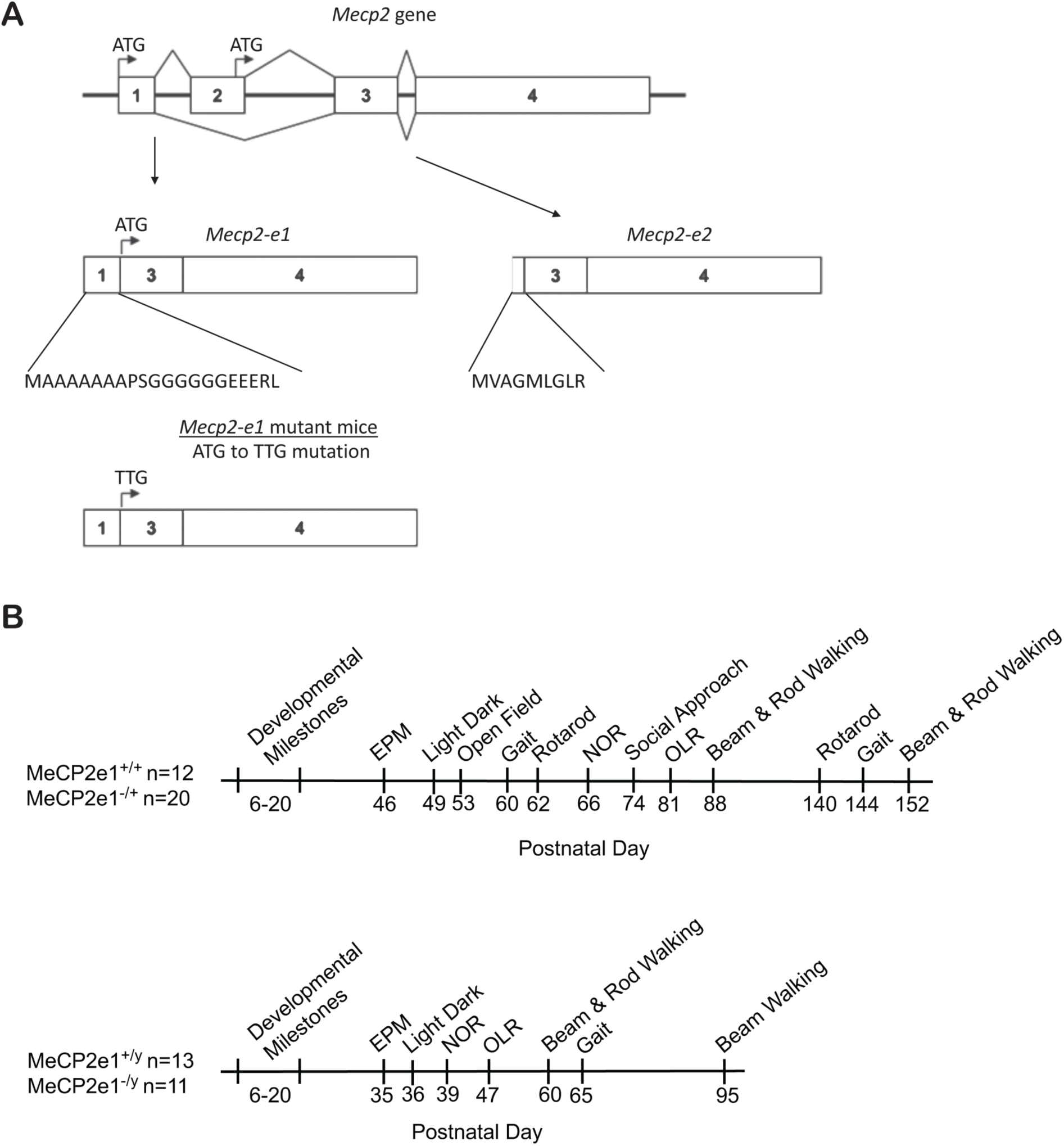
MeCP2-e1 isoform splicing and experimental design. **A.** The *Mecp2* transcript is alternatively spliced into the e1 and e1 isoforms with different N-terminals (amino acid sequence listed). MeCP2-e1 mutant mice have a single nucleotide substitution that results in a loss of the start codon (ATG to TTG mutation) and no translation of the MeCP2-e1 protein. The MeCP2-e2 isoform is still transcribed and translated. **B.** The timeline for behavioral analysis of MeCP2e1 mutant females (*Mecp2e1^-/+^*) and wildtype littermates (MeCP2e1^+/+^) and mutant males (MeCP2e1^-/y^) and wildtype littermates (MeCP2e1^+/y^).

The majority of previous work on MeCP2 mouse models of RTT has focused on *Mecp2^-/y^* hemizygous male mice as they show early and progressive symptom onset and lethality. Previous work on female mouse models of RTT has focused primarily on heterozygous *Mecp2* mutations in 20–40 percent of all cells (7, 8). Initial characterization of heterozygous knockout females (*Mecp2^tm1.1bird-/+^*) reported delayed phenotype development until four to six months of age (9, 10). Recent, more detailed work in female heterozygous mouse models has revealed mixed findings for the age of phenotype development, depending on background strain and mutation type (11–16). However, to date there has been limited exploration of the time course of phenotype development and no behavioral characterization of isoform specific manipulations of MeCP2 in female mouse models. Given that RTT occurs almost exclusively in females and there is no current effective treatment, there is great need to develop a preclinical female mouse model of RTT that recapitulates the delayed behavioral phenotypes observed in human females with RTT.

While MeCP2 is most highly expressed in neurons, human female RTT patients exhibit immune, microbiome, mitochondrial, and metabolic manifestations that are likely reactionary to the causal mutation and neuronal dysfunction (17–25). Metabolic dysfunctions are observed in RTT patients and mouse models, but whether these are consequences of inflammation or more directly mediated by *MECP2* mutation is currently unknown. While RTT girls are generally not obese, perhaps due to the inability to self-feed, they have elevated morning leptin and adiponectin serum levels (17). Furthermore, RTT has significant commonalities with mitochondrial disorders, including elevated lactate and pyruvate in blood and cerebral spinal fluid (26), as well as markers of oxidative damage (27) that are also seen in mouse models (28, 29). Lipid metabolism was identified in an unbiased screen for rescue of RTT mouse phenotypes (30) and was later shown to result in metabolic alterations in RTT mice (31). Metabolic or oxidative stress alterations have been modified in several therapeutic trials of RTT mouse models (20), with some discrepancies by strain and age (32, 33). Since the most promising therapeutics for RTT target metabolic pathways (34), such as AKT/mTOR (35, 36), oxidative stress (28, 37), and lipid metabolism (30, 38), it is imperative to characterize developmental, metabolic, and motor phenotypes in female RTT mouse models.

To address the role of MeCP2-e1 in the development of phenotypes relevant to RTT, we performed a longitudinal assessment of *Mecp2e1^-/+^* female compared to *Mecp2e1^-/y^* male mice across development and into adulthood. We assessed developmental milestones during the first three weeks of life and monitored the progressive development of hind limb clasping and changes in body weight into adulthood. In addition, we performed assays examining anxiety, sociability, motor function, cognition, and metabolism. A delayed and progressive onset of hind limb clasping, increased body weight, and motor impairments was observed, reminiscent of the delayed symptom onset observed in females with RTT. Our findings indicate that changes in body weight may exacerbate motor impairments in the *Mecp2e1^-/+^* female mice, suggesting that metabolic phenotypes may serve as a novel therapeutic avenue for RTT.

## Results

### Both female and male MeCP2-e1 mutant mice show an onset of hindlimb clasping phenotypes by two months that precede a progressive increase in body weight

To examine potential early phenotypes in *Mecp2e1* deficient mice, a panel of developmental milestones was assessed across the first 21 days of life. The majority of measures showed no significant differences in mutant development compared to sex matched wild-type littermates (Supplemental Table S1). The two exceptions were slight delays in pinnae detachment in *Mecp2e1^-/+^* females and cliff avoidance in *Mecp2e1^-/y^* males. However, all other measures were indistinguishable from wild-type, indicating normal pre-weaning development in mutant animals.

Post weaning body weight and hind limb clasping scores were assessed weekly and showed progressive symptom development in both male and female mutants (Figure 2 and Supplemental Table S2). Female *Mecp2e1^-/+^* mice showed clasping scores that were significantly different from wild-type beginning at PND 52 that progressed, then stabilized after PND 69. This neurologically-based hind limb clasping phenotype in *Mecp2e1^-/+^* females preceded a progressive significant increase in body weight starting at PND 74. *Mecp2e1^-/y^* male mice showed a similar but slightly delayed onset of significant differences from wild-type in hindlimb clasping by PND 61 and significantly higher body weights beginning at PND 92. The hind limb clasping phenotype was more severe in male compared to female mutants, but increases in body weight relative to wild-type littermates were similar in magnitude between sexes.

**Figure 2.**
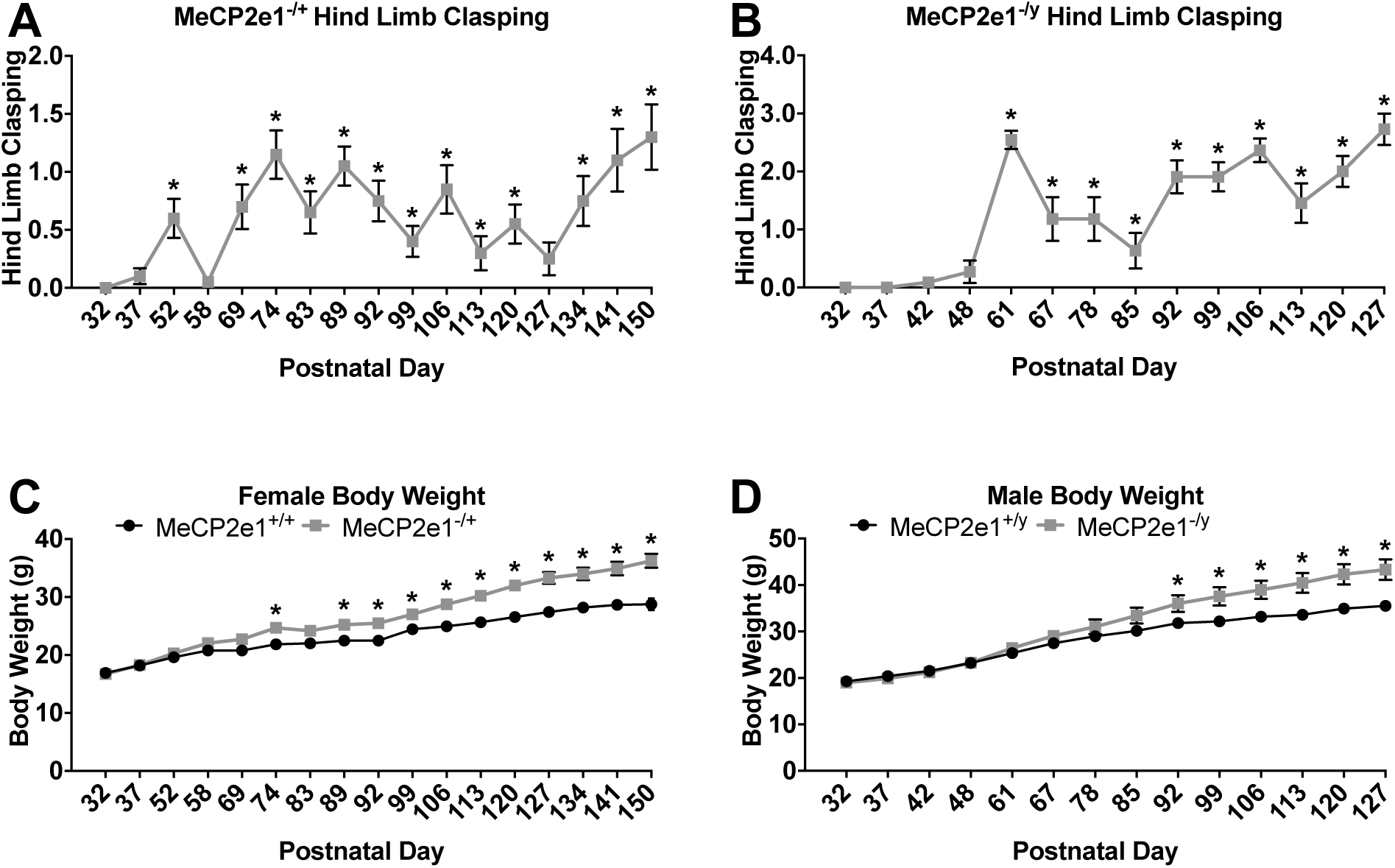
Symptom development across age in male and female MeCP2e1 mutant mice. Hind limb clasping was assessed weekly using a rating scale from 0 (no clasping) to 5 (severe). Data were analyzed using an exact binomial test that tests whether there were significantly more mutant animals with clasping scores > 0 for each time point (PND). P-values were Benjamini-Hochberg corrected across time points. **A.** Hind limb clasping in females. **B**. Hind limb clasping in males. Body weight (grams) was taken weekly across development and analyzed by linear mixed effects model (see methods) and Benjamini-Hochberg corrected comparisons were conducted between genotypes for each PND. **C.** Body weight in females. **D.** Body weight in males. * mutant versus wildtype corrected p value < 0.05, statistical tests shown in detail in Supplemental Table S2. Mean +/- standard error of the mean. PND: postnatal day.

Both female and male mutant mice (PND 46 and 53) showed increased time in the open arm of the elevated plus-maze (Figures S1 and S2) as previously found for MeCP2e1^-/y^ males (39) and *Mecp2^tm1.1bird-/+^* females (16) at similar ages. There were no significant differences between genotypes for the number of transitions between arms indicating the increased time in the open arm was not due to changes in motor function. Female *Mecp2e1^-/+^* mice did not differ from wild-type littermates in the light – dark exploration task (Figure S1), however *Mecp2e1^-/y^* males did spend significantly less time in the dark chamber than wild-type littermates with no difference in the number of transitions (Figure S2). The inconsistent findings on the two measures of anxiety indicate potential abnormalities in more subtle aspects of the two tasks and are consistent with previously observed discrepancies between phenotypes on the two tasks for *Mecp2^tm1.1bird-/+^* females (16). There were no significant differences between mutant and wild-type females in the open field text of exploratory behaviors except in the measure of center time, where *Mecp2e1^-/+^* mice spent significantly less time in the center of the arena than wild-type littermates (Figure S3). Adding body weight as a covariate did not alter the results in these tasks (Table S3 and S4).

### Female MeCP2-e1 mutant mice exhibit early motor deficits impacted by age-related body weight differences

Motor function was assessed using several different tasks including analysis of gait, beam and rod walking latency, and accelerating rotarod performance. Female *Mecp2e1^-/+^* mice showed significant impairments on several measures of gait performance (Figure 3). At PND 60 mutant females showed significant impairments in stride length, hind base, and paw separation. However, by PND 144 only the differences in stride length were significant. Body weight was a significant covariate for the measure of stride length but did not alter the significance of the effect of genotype (Table S3). Adding body weight as a covariate did alter the main effect of genotype for the measure of hind base, indicating that this measure of gait may be more sensitive to weight changes. *Mecp2e1^-/y^* males did not show any significant difference from wild-type littermates on any of the gait measures (Figure S4 and Table S4).

**Figure 3.**
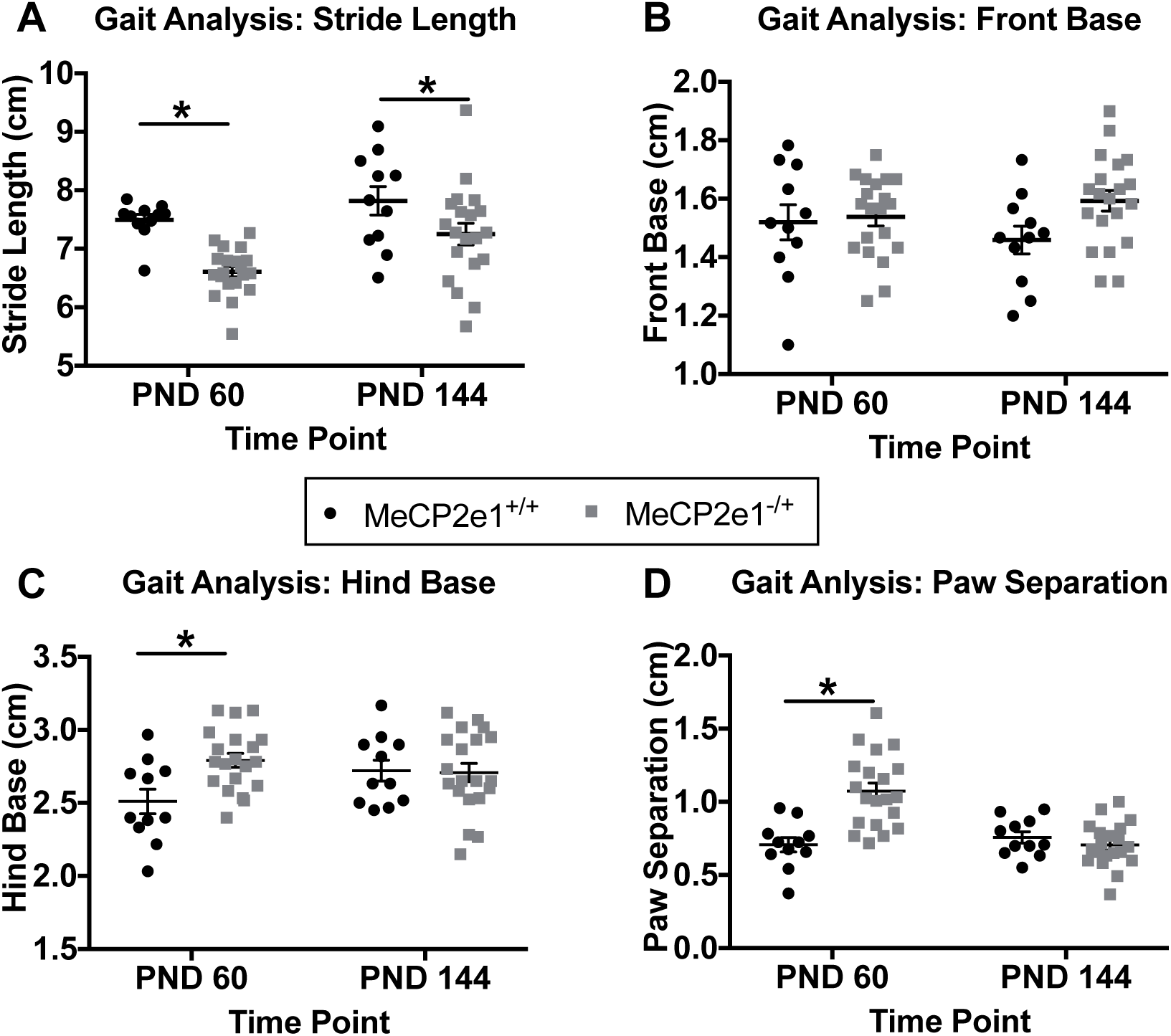
Female mutant mice show impairments in gait in early development. **A.***Mecp2e1^-/+^* mice show decreased stride length at both PND 60 and 144. Impairments measures of front base (**B**), hind base (**C**) and paw separation (**D**) only were significantly altered at PND 60. Mean +/- Standard error of the mean. * p<0.05 Benjamini-Hochberg corrected comparisons between genotypes, statistical tests shown in detail in Supplemental Table S3.

Female *Mecp2e1^-/+^* mice showed significant impairments in beam walking at both PND 88 and 152 (Figure 4 and Table S3). Deficits were specific to the smallest diameter beam (beam 3) at the younger time point and to the widest diameter beam (beam 1) at the older time point, potentially representing impairments in different aspects of the task with symptom progression. This task was also sensitive to alterations in body weight. When added as a covariate, the main effect of genotype was no longer significant at either time point, indicating that increased body weight in the mutant females is a significant contributor to impairments in this task. In comparison, female mutant mice did not show any significant impairment on rod walking (Figure S5 and Table S3). Female *Mecp2e1^-/+^* mice did show progressive impairments in the accelerating rotarod (Figure 4 and Table S3). At PND 62, performance between wild-type and mutant females was equivalent across all three days of training, indicating normal motor performance and learning. At PND 140, *Mecp2e1^-/+^* mice showed significant impairments across all three days of training. The impairments at PND140 were sensitive to changes in body weight. When body weight was included as a covariate, the main effect of genotype was no longer significant indicating that at least a portion of the impairment in performance was due to increased weight in the mutants. Motor and phenotypic symptoms were also delayed relative to those observed in heterozygous whole exon deletion female models (16), suggesting that some MeCP2e-1 roles in motor skills may be compensated for by MeCP2-e2.

**Figure 4.**
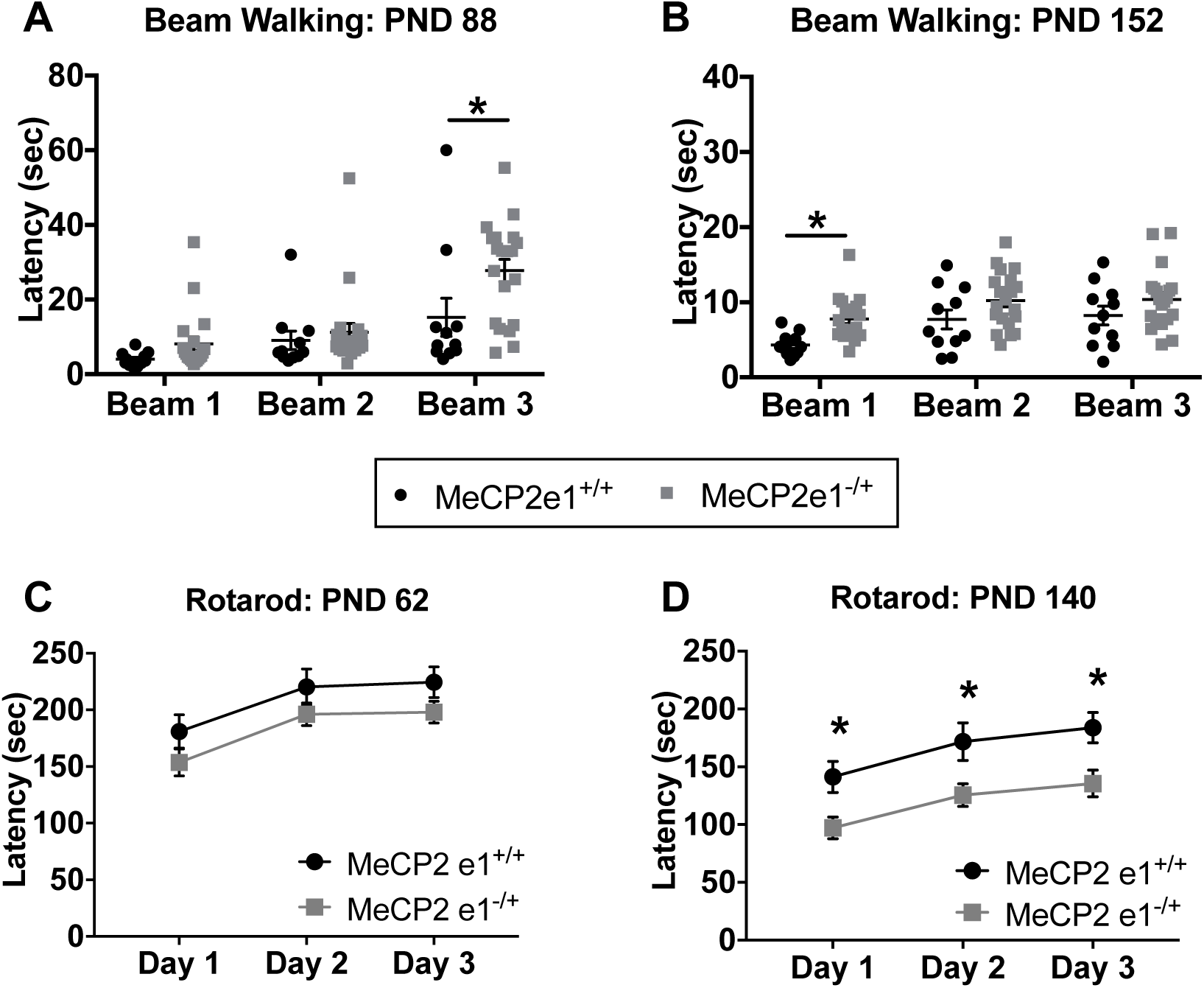
Female mutant mice show impairments in motor tasks. **A.** *Mecp2e1^-/+^* mice show increased latency to transverse beam 3 (smallest diameter) at PND 88. **B.** *Mecp2e1^-/+^* mice show increased latency to transverse beam 1 (largest diameter) at PND 152. **C.** *Mecp2e1^-/+^* mice show similar performance to wildtype littermates on the accelerating rotarod across three days of training at PND 62. **D.** *Mecp2e1^-/+^* mice show a decreased ability to remain on the accelerating rotarod (shorter latency) compared to wildtype littermates across three days of training at PND 140. Mean +/- standard error of the mean. * p<0.05 Benjamini-Hochberg corrected comparisons between genotypes, statistical tests shown in detail in Supplemental Table S3.

Male *Mecp2-e1^-/y^* mice also showed significant impairments on the beam walking but not rod walking task (Figure S6 and Table S4). At PND 60, *Mecp2e1^-/y^* mice showed significant impairments relative to wild-type littermates on the smallest diameter beam (beam 3). At PND 95, *Mecp2e1^-/y^* mice were significantly impaired on all three diameter beams. At the PND 60 but not PND95, adding body weight as a covariate resulted in a loss of the significant genotype effect. This suggests that the impairments at the earlier time point may be driven by body weight but that later impairments are largely motor in origin.

### Lack of significant deficits in sociability or short-term memory in young female MeCP2-e1 mutant mice

Male *Mecp2e1^-/y^* mice were previously shown to have deficits in social behavior (5) and consequently, we examined sociability of female *Mecp2e1^-/+^* mice in the three chambered social approach task (Figure S7). *Mecp2e1-/+* mice performed similarly to wild-type littermates in the number of transitions between chambers during both the habituation and testing phases, indicating that any motor impairments minimally impacted the task. Both *Mecp2e1^-/+^* mice and wild type littermates showed significant preferences for the novel mouse over the novel object both in chamber time and sniff time. Together this suggests that female *Mecp2e1^-/+^* mice do not have impairments in sociability at PND 74.

Both male and female mutant mice were examined for cognitive performance using the short-term memory version of the novel object task and novel location task (40). Both *Mecp2e1^-/+^* and wild-type female mice showed a significant preference for the novel object over the familiar object and there were no differences in total exploration time or distance traveled at training or testing (Figure S8 and Table S3). For the object location version of the task, *Mecp2e1^-/+^* mice showed a significant preference for the object in the new location and wild-type littermates showed a marginally significant (p<0.06) preference. There was no difference between genotypes for total exploration at training or testing and no difference in total distance traveled. Together, these findings indicate intact short-term memory for both novel objects and object locations in *Mecp2e1^-/+^* mice. Similarly, both male wild-type and *Mecp2e1^-/y^* mice showed a significant preference for the novel compared to familiar object during novel object recognition testing (Figures S8 and Table S4). However, *Mecp2e1^-/y^* mice did show significantly lower overall exploration during training and higher exploration during testing compared to wild-type littermates. For the object location version of the task, wild-type males showed a significant preference for the object in the new location and *Mecp2e1^-/y^* mice showed a marginally significant preference (p<0.06). Again, *Mecp2e1^-/y^* mice showed higher overall exploration of the objects during training but not during testing. Overall, these findings suggest intact learning and short-term memory in both male and female mutant mice.

### MeCP2e1 mutant males exhibit significantly increased body fat despite reduced food intake due to genotype effects on energy expenditure

Since previous studies suggest metabolic defects occur in both RTT females and *Mecp2*/MeCP2 deficient mouse models (20, 28, 31, 41) and because of the progressive elevated body weight in MeCP2-e1 mutant mice, we sought to better characterize the metabolic phenotypes at an early stage of disease progression (PND 53–70 in males, PND 86–91 in females) when no significant differences in body weight were observed by ANOVA (Figure S10). Dual-energy, X-ray absorptiometry (DEXA) was performed to measure fat mass and bone mineral density and area. DEXA scans revealed that both *Mecp2e1^-/y^* males and *Mecp2e1^-/+^* females had elevated percent fat (Figure 5A, 5B) and elevated fat mass (Figure 5C, 5D). *Mecp2e1^-/y^* mice had elevated percent fat and fat mass compared to wild-type male littermates, and *Mecp2e1^-/+^* mice had elevated percent fat and fat mass compared to wild-type female littermates. These differences in *Mecp2e1* deficient mice were surprising as weights were similar between *Mecp2e1^-/y^* and controls (Figure 5E) as well between *Mecp2e1^-/+^* and control females (Figure 5 F). The results are consistent with genotype altering nutrient partitioning and inducing a shift in body composition independent of changes in body weight. DEXA analysis also revealed that *Mecp2e1-/y* males but not *Mecp2e1^-/+^* females had reductions in bone mineral content (Figure S12A-B), bone area (Figure S12 C-D) and body length (Figure S12E-F).

**Figure 5.**
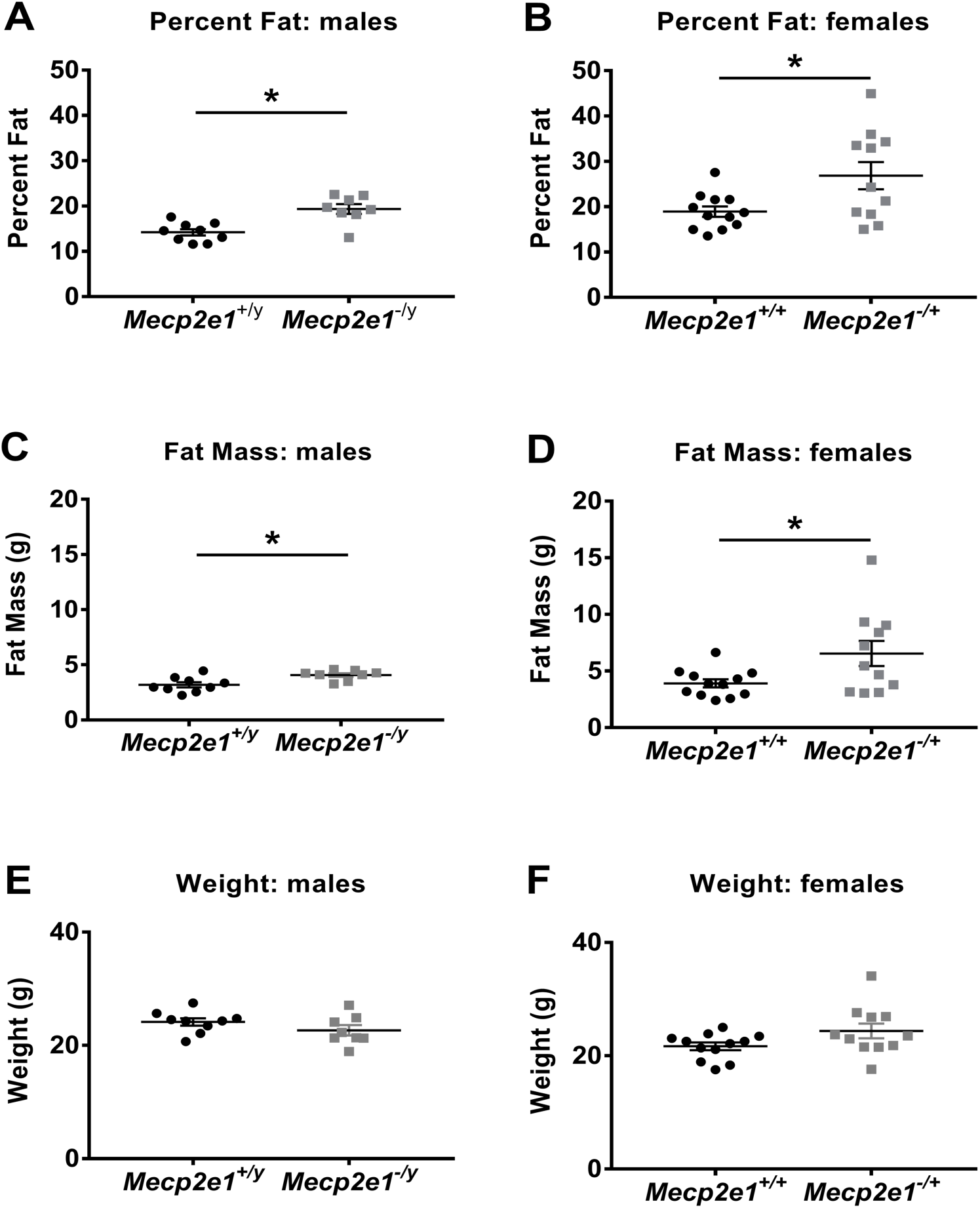
DEXA analysis reveals body composition alterations for *Mecp2e1* mutant mice. For these studies *Mecp2e1^+/y^* (N=9), *Mecp2e1^-/y^* (N=8), *Mecp2e1^-/+^* (N=11), and *Mecp2e1^+/+^* (N=12) were analyzed. **A.** A significant (p=0.00096) elevation for percent fat in *Mecp2e1^-/y^* compared to control *Mecp2e1^-/+^* mice is shown. **B.** A significant (p=0.0183) elevation in percent fat for *Mecp2e1^-/+^* compared to control *Mecp2e1^+/+^* littermates. **C.** Fat mass in grams is elevated in *Mecp2e1^-/y^* (p=0.028) and **D.** *Mecp2e1^-/+^* (p=0.0077) compared to *Mecp2e1^+/y^* and *Mecp2e1^+/+^* control littermates respectively. **E.** Weights are comparable between *Mecp2e1^+/y^* and *Mecp2e1^-/y^* mice (p=0.2). **F** *Mecp2e1^+/+^* and *Mecp2e1^-/+^* mice have similar weights at the time of DEXA (p=0.07). P values were computed using an unpaired, two-tailed student’s t-test. * = p value < 0.050. Error bars correspond to Standard Error of the Mean (SEM).

To identify potential causes of the increased fat mass and progressive increase in body weights, food intake, energy expenditure, respiratory exchange ratio, and physical activity of *Mecp2e1* deficient mice were measured using a Comprehensive Lab Animal Monitoring System (CLAMS). Interestingly, despite having increased fat, *Mecp2e1^-/y^* mice consumed significantly less calories than *Mecp2e1+/y* wild-type male littermates (Figure 6A) and this was driven primarily by significantly reduced consumption during the light phase (Figure 6C). In contrast *Mecp2e1^-/+^* females did not show significant effects on food consumption overall or during the light phase (Figure 6B, 6D). Food consumption in the dark phase was not significantly different between *Mecp2e1^+/y^* and *Mecp2e1^-/y^* mice (Figure S13A) nor between *Mecp2e1^+/+^* and *Mecp2e1^-/+^* females (Figure S13B).

**Figure 6.**
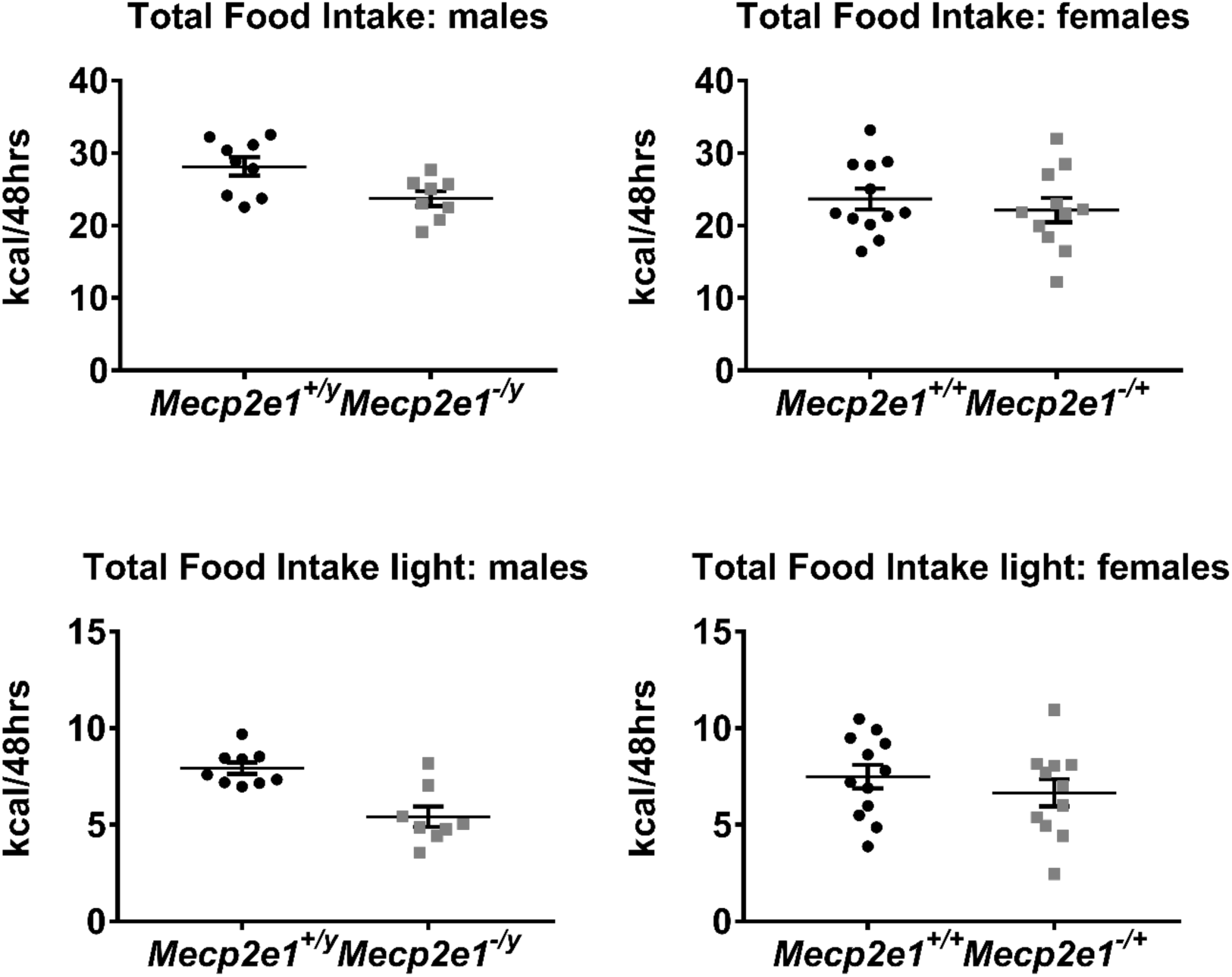
*Mecp2e1^-/y^* mice consume less calories during the light phase. **A.** A significant (p=0.0173) reduction for total cumulative food intake for 48 hours represented as kilocalories per 48 hours (kcal/48hr) between *Mecp2e1^+/y^* control and *Mecp2e1^-/y^* groups is shown. **B.** Total cumulative food intake is comparable between *Mecp2e1^-/+^* and *Mecp2e1^+/+^* control littermates. **C.** Sum food intake during the light phase reveals a significant reduction (p=0.0007) in *Mecp2e1^-/y^* compared to *Mecp2e1^+/y^* littermate controls. **D.** Sum food intake in the light phase is similar in *Mecp2e1^-/+^* and *Mecp2e1^+/+^* control littermates. For these studies *Mecp2e1^+/y^* (N=9), *Mecp2e1^-/y^* (N=8), *Mecp2e1^-/+^* (N=11), and *Mecp2e1^+/+^* (N=12) were analyzed. P values were computed using an unpaired, two-tailed student’s t-test. * p < 0.05.

To determine the relationship between metabolism and physical activity, the total number of X-axis and X-axis ambulatory beam breaks were compared across *Mecp2e1* genotypes (Figure 7). *Mecp2e1^-/y^* males showed reduced locomotion compared to wild-type littermates as shown by significant reductions in X-axis beam breaks (Figure 7A) and X-axis ambulation (Figure 7C). *Mecp2e1^-/+^* females showed a trend towards lower levels of activity, although the effect was not significant (Figure 7B, 7D). The overall reduction in *Mecp2e1^-/y^* activity was due to reduced X-axis total activity and X-axis ambulation during the dark cycle (Figure S14A, S14 C). In contrast *Mecp2e1^-/+^* females had normal X-axis total activity and X-axis ambulation during the dark cycle (Figure S14B, S14D).

**Figure 7.**
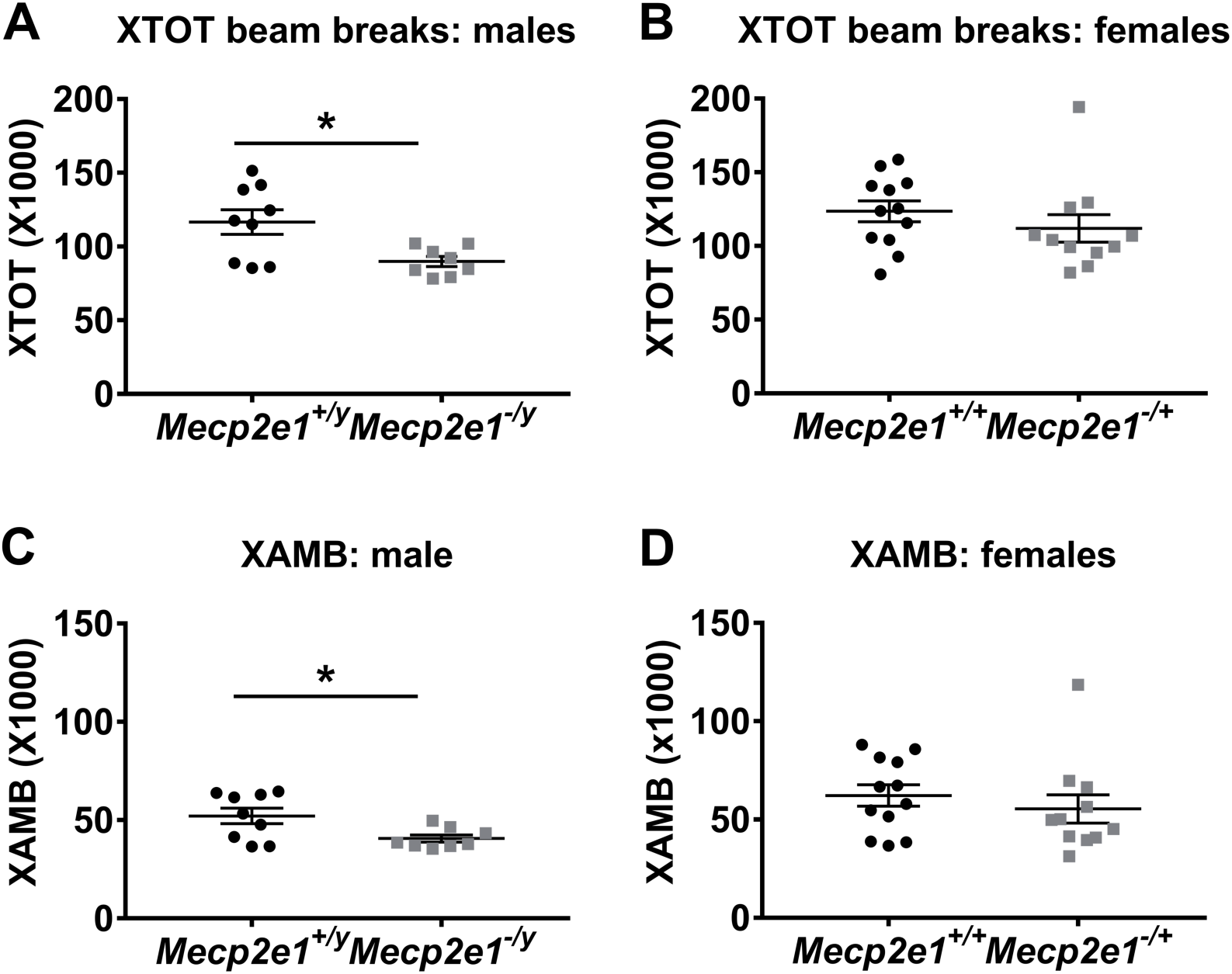
***Mecp2e1^-/y^* mice show reduced locomotion. A.** A significant (p=0.023) reduction for Total 48 hour (h) cumulative X beam breaks between *Mecp2e1^+/y^* control and *Mecp2e1^-/y^* groups is shown. **B** Total 48ho cumulative X beam breaks show a trend toward reduction in *Mecp2e1^-/+^* compared to *Mecp2e1^+/+^* controls. **C** Total 48h cumulative X beam break analysis reveals a significant reduction (p=0.013) in *Mecp2e1^-/y^* mice compared to *Mecp2e1^+/y^* control littermates. **D** A trend toward reduction in total 48h X ambulation in *Mecp2e1^-/+^* mice compared to *Mecp2e1^+/+^* controls is observed. For these studies *Mecp2e1^+/y^* (N=9), *Mecp2e1^-/y^* (N=8), *Mecp2e1^-/+^* (N=11), and *Mecp2e1^+/+^* (N=12) were analyzed. P values were computed using an unpaired, two-tailed student’s t-test. * p < 0.05.

Respiratory exchange ratio (RER) and energy expenditure were also examined as a possible explanation for the increased fat and body weights of MeCP2-e1 mutant mice. *Mecp2e1^-/y^* mice exhibited significantly increased RER (Figure 8A), representing a shift towards increased carbohydrate oxidation compared to wild-type male littermates. Energy expenditure was also significantly lower in *Mecp2e1-/y* mice compared to control males (Figure 8C), while *Mecp2e1^-/+^* females were not significantly different than female controls by either measurement (Figure 8B, 8D). Analysis of covariance (ANCOVA) normalizing for either body weight (p= 0.006482) or lean mass (p= 0.01526) confirmed the reduction energy expenditure in *Mecp2e1^-/y^* males. In contrast, energy expenditure did not differ in *Mecp2e1^-/+^* females with either body weight (p= 0.2257) or lean mass (p= 0.5225) normalization by ANCOVA (Supplemental Table S5). Interestingly, the alterations in RER and energy expenditure in *Mecp2e1^-/y^* males were primarily observed in the dark phase (6PM – 6AM) of the circadian cycle (Figure S15A,15 C), although RER was not altered in the light phase when energy expenditure was also reduced (Figure S16A, S16 C). *Mecp2e1^-/+^* females did not differ from controls in RER and energy expenditure during the dark phase (Figure S15B, S15D) as well as the light phase (Figure S16B, S16D). Together, these results demonstrate a significant genotype effect of fuel usage and energy expenditure in the active dark hours that, together with reduced activity, appear to predispose MeCP2-e1 deficient male mice to lipid accumulation and weight gain despite reduced food intake.

**Figure 8.**
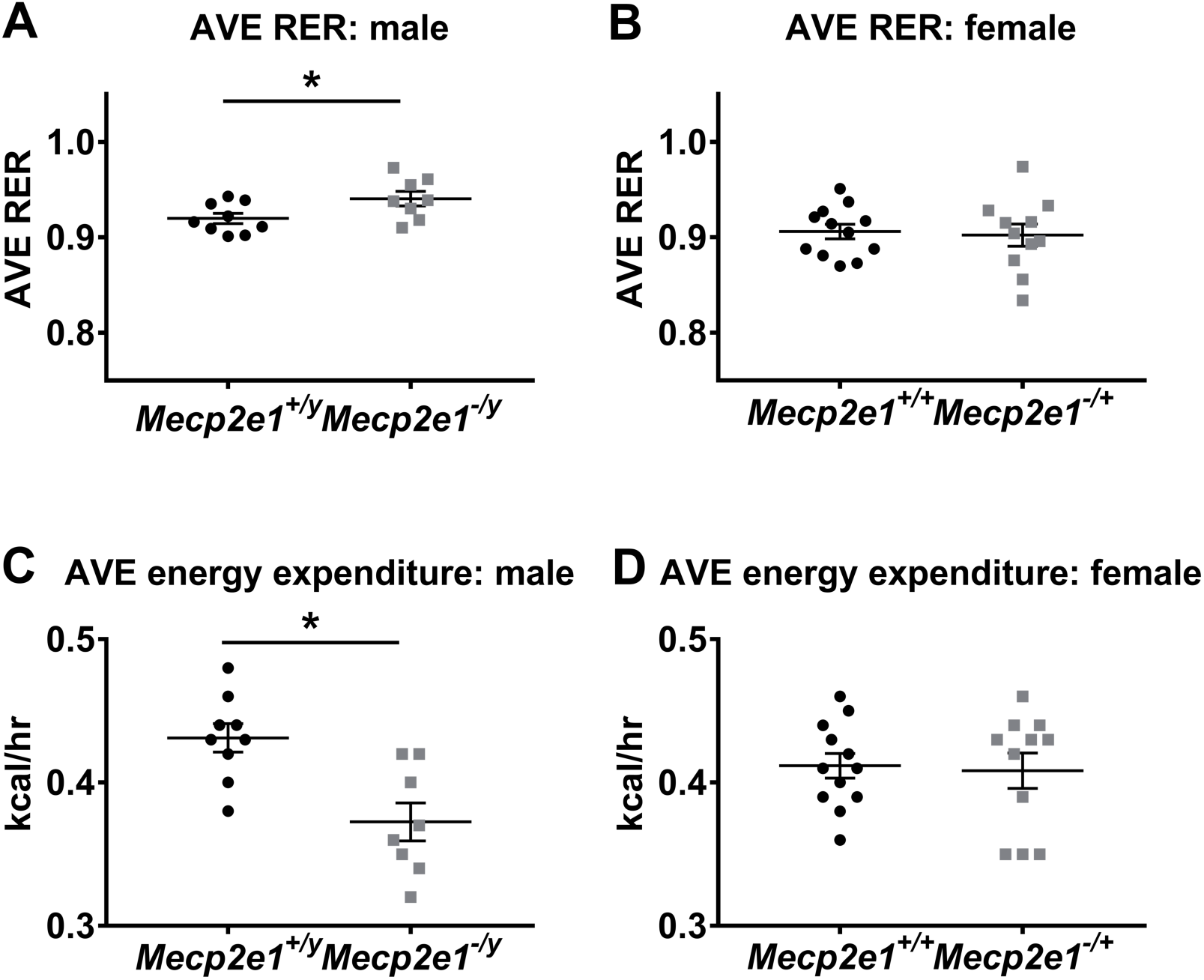
*Mecp2e1^-/y^* mice have elevated RER and reduced energy expenditure. A. Average respiratory exchange ratios (AVE RER) shows elevation (p=0.0368) in *Mecp2e1^-/y^* (N=8) compared to littermate controls *Mecp2e1^+/y^* (N=9). B. Average RER is similar between *Mecp2e1^-/+^* (N=11) and *Mecp2e1^+/+^* (N=12) control littermates. C. Average energy expenditure (AVE Heat) for *Mecp2e1^-/y^* mice (N=8) is significantly reduced (p=0.00194) compared to controls *Mec2pe1^+/y^* (N=9). D. Average energy expenditure was similar between *Mecp2e1^+/+^* control (N=12) and *Mec2pe1^-/+^* (N=11) mice. P values were computed using an unpaired, two-tailed student’s t-test. * p < 0.05.

### MeCP2-e1 mutant female profiling of liver lipids demonstrates a distinct profile of triglycerides

Previous analysis of MeCP2 null *Mecp2^-/y^* liver has shown grossly elevated accumulation of triglyceride and cholesterol compared to control liver (31). Therefore, as *Mecp2e1^-/+^* females showed significant increases in fat mass compared to controls we explored possible changes in lipid accumulation in livers from mutant females. Untargeted lipid profiling by mass spectrographic analysis was performed on *Mecp2e1^-/+^* liver samples following CLAMS and DEXA analysis of the same animals. Relative lipid abundance from liver was determined by peak height compared to standards and analyzed using MetaboAnalyst 4.0 statistical analysis package. Figure 9A demonstrates that *Mecp2e1^-/+^* liver samples could be separated from wildtype female control liver samples in a partial least squares discriminant analysis (PLS-DA) of lipid abundance. Several individual lipids were significantly increased by raw p values, although none were significant after FDR correction (Figure 9B). Of the top lipid differences ranked by raw p values, triglycerides 547, 542, 565 and 512 were elevated (Figure 9C). Together, these results suggest that *Mecp2e1^-/+^* female mice have elevated triglycerides compared to their wild-type littermates.

**Figure 9.**
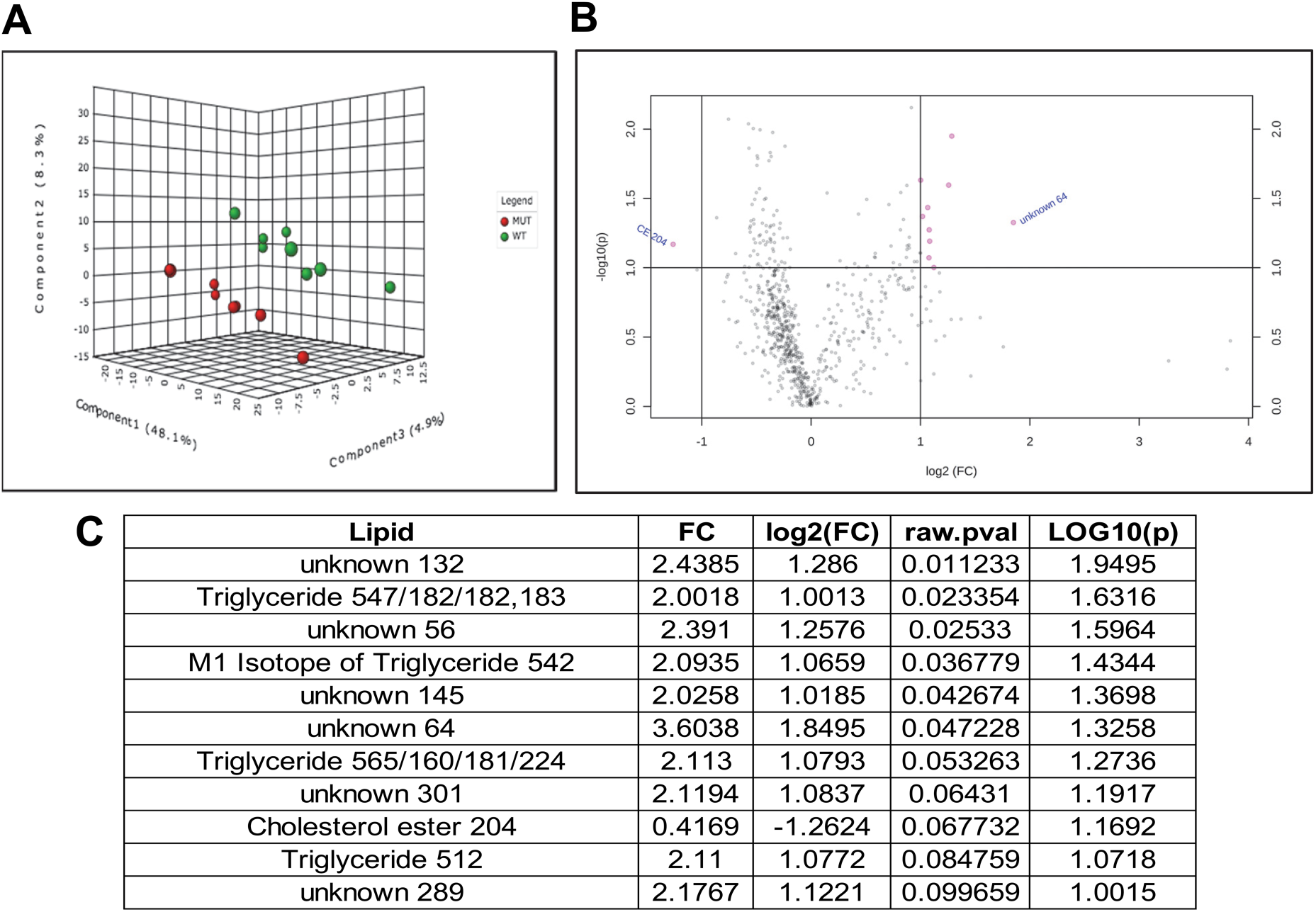
Untargeted lipid profiling analyses of *Mecp2e1^-/+^* liver indicate elevated triglycerides. **A.** Partial least squares discriminant analysis (**PLS-DA**) shows a clear separation between *Mecp2e1^-/+^* samples (MUT, green) and control littermate (WT, red) female liver samples based on untargeted lipid profiling. **B.** A Volcano plot of fold change (x axis) and raw p values of genotype differences based on lipid abundance determined by mass spectroscopic analysis. Pink circles represent lipids with a log2 fold change >1.0 and a top raw p values rank, with the identities of the know lipids shown in **C**. Of the known lipids, triglycerides were the most represented category of lipids that had elevated abundance in *Mecp2e1^-/+^* liver samples compared to control liver samples.

## Discussion

Mutations in *MECP2* cause the majority of RTT cases and despite many years of research we still have a limited understanding of how MeCP2 causes a regressive disease phenotype in heterozygous and mosaic females. Furthermore, there has been limited progress on how each isoform of MeCP2 (MeCP2-e1 and -e2) contributes to the diverse functions of MeCP2. This study provides some important novel findings about the specific role of MeCP2-e1 deficiency on disease progression, with a specific focus on the female phenotypes that are expected to be most important to understanding the complex pathogenesis of human RTT. First, we demonstrate that female MeCP2-e1 deficient mice show hind-clasping phenotypes at the same stage as male MeCP2-e1 deficient littermates, around 2 months of age. Second, both male and female MeCP2-e1 deficient mice exhibit significant and progressive weight gain, but at a time point later than significant differences in hindlimb clasping, fat mass, and metabolism were observed. Third, female MeCP2-e1 deficient mice exhibit significant motor deficits at early stages of disease progression which are influenced by body weight differences. Lastly, we demonstrated significant *Mecp2e1* genotype effects on measurements of fat mass, bone mineral content, food intake, physical activity, and energy expenditure in mice prior to the onset of weight gain, suggesting that metabolic disruptions due to MeCP2-e1 deficiency occur early in disease progression. Together, these results demonstrate a timeline of RTT-relevant phenotype progression in a mouse model of an actual RTT mutation that should be important for better understanding and treatment of human RTT.

Because of the complex MeCP2 mutant/wild-type mosaicism observed in *Mecp2* mutant female mice, the majority of RTT pre-clinical therapy trials in mice are performed only on *Mecp2* null male mice lacking MeCP2 in all cells (42–47). Reasons cited for only testing *Mecp2* null males include the obvious phenotype (death by 8–12 weeks) and the lack of confounding X chromosome inactivation mosaicism. However, *MECP2* null mutations are not seen in human RTT patients, suggesting that *Mecp2* mutant mice designed from RTT-causing human mutations are more construct-relevant models (5, 48–50). A variety of genetic models have been developed including whole exon deletions (9, 10), specific DNA binding domain deletions (14), and a variety of knock-in point and truncation mutations that model human mutations (39, 51–55). The majority of these models develop deficits similar to those observed in RTT patients, including abnormal breathing, gait abnormalities, tremors, EEG abnormalities, and in some cases cognitive deficits, social abnormalities, and changes in anxiety behaviors (56). However, the complete loss of MeCP2 expression and the rapid and severe deterioration in the male models do not capture the delayed onset of symptoms and mosaic inactivation patterns of *MECP2* expression in RTT females. While death is the primary phenotype in male RTT mouse models, motor, respiratory, metabolic, and severe gastrointestinal symptoms are more medically relevant to the variable range of phenotypes in female RTT patients, since >70% of RTT women are still alive by age 45^35^. Our results demonstrating significant phenotypes of hindlimb clasping, increased fat mass, and motor deficits by two months of age in female MeCP2-e1 deficient mice therefore provide an important advancement to the existing literature.

*Mecp2e1^-/+^* females developed abnormal behaviors by 6–7 weeks of age, similar to previous observations in *Mecp2^-/+^* mutant females heterozygous for exon 3–4 deletion affecting both *Mecp2* isoforms (11, 16, 57). Motor deficits are among the most common and consistently observed phenotypes in male and female mouse (12, 16, 42, 58–62) and rat (63–65) models of RTT. Both *Mecp2e1^-/+^* females and males develop progressive motor impairments over time. Female *Mecp2e1^-/+^* mutants show early motor impairments (8.6 weeks of age) in gait but not in rotarod. By later timepoints (ages 12–20 weeks) the motor phenotypes became fully evident in the mutant females in gait, beam walking and rotarod impairments. The development of full motor phenotypes is delayed relative to *Mecp2bird^-/+^* females tested at the same ages in our previous study (16) and more heavily influenced by body weight (Supplemental Figure 11). The increased body weight in *Mecp2e1^-/+^* females was a significant covariate or influenced the main effect of genotype on the majority of motor measures, indicating that the metabolic phenotypes in this model may exacerbate motor impairments.

Recent identification of mutations that enhance survival of *Mecp2^-/y^* males has established the link between *Mecp2* and lipid metabolism (30–32). Consistent with these findings our results show that *Mecp2e1^-/+^* females at 12 to 14 weeks of age and *Mecp2e1^-/y^* males at 7 to 11 weeks of age display significant elevation in body fat percent and fat mass without significant elevation in weight on a C57BL6/J background. Interestingly, results most relevant to our studies show that *Mecp2*^-/y^ mice on a 129 background are significantly heavier beginning at 7 weeks of age (31). Furthermore, while Kyle et al showed reduced RER in *Mecp2^-/y^* mice in both the light and dark cycle (31) our results show that *Mecp2e1^-/y^* mice have elevated RER predominantly in the dark cycle. In terms of energy expenditure, we show that *Mecp2e1^-/y^* males have reduced dark and light phases energy expenditure even after adjusting for body weight or lean mass, consistent with reduced heat production during the light cycle for *Mecp2^-/y^* null mice on a 129 background as shown previously (31). The genetic background or genetic mutation difference of our *Mecp2e1^-/y^* mice on a C57BL6/J background may explain differences between studies, although the studies are consistent in showing metabolic alterations including decreased energy expenditure and increased mass in RTT mouse models. Bone defects in *Mecp2* null males were previously observed in *Mecp2* null mice (66), which is consistent with our findings in MeCP2-e1 deficient mice.

In support of a possible decreased energy expenditure in RTT mouse models across genetic backgrounds, Previous metabolic analyses of RTT model mice carrying the B6.129P2(C)-Mecp2tm1.1Bird/J or “Bird” null allele have shown that *Mecp2*^-/+^ females have elevated body weight on 129 or C57BL6/J/129 mixed backgrounds (9, 10, 16, 51, 53). Similarly, genetic background clearly affects weight in *Mecp2* null (Bird) male mice as they were underweight on a C57BL6/J background while overweight on a 129 background (9, 16) or mixed C57BL6/J/129 background (67). Interestingly Kerr et al also showed elevated weight in mice bearing the floxed, undeleted *Mecp2* (Bird) allele on a C57BL6/J/129 mixed background, suggesting an effect of *Mecp2* 3’UTR on metabolic phenotype (68). Recently, Gigli et al showed that *Mecp2^-/y^* male mice bred on a CD1 background have reduced weight at 6 weeks followed by significantly elevated adult weight, while CD1 *Mecp2^-/+^* females exhibit elevated weight starting at 17 weeks compared to wild-type littermates (69). Consistent with these findings, Torres-Andrade showed that *Mecp2* null (Bird) males on a mixed C57BL6/J/129 background were overweight by 7 weeks of age due to increased fat mass and that this was not due to increased food intake (70), consistent with our results.

Conditional knockout experiments also have revealed common body weight effects from *Mecp2* deletion in specific neuronal subtypes. For example, *Mecp2* deletion in forebrain neurons led to significant weight gain compared to littermate controls at 13 weeks in the C57BL6/J background (71). Conditional *Mecp2* (Bird allele) knockout in hypothalamic neurons resulted in weight gain due to fat mass expansion in *Mecp2^-/y^* males by 7 weeks on a 129/FVB background (72). Also, *Mecp2^-/y^* males with a conditional deletion in only excitatory neurons became overweight by 6 weeks of age on a 129/FVB background (73). Conditional knockout of *Mecp2* in POMC neurons resulted in increased body weight at 16 weeks in *Mecp2^-/y^* (Bird) males on a C57BL6/J background (74). Cumulatively these experiments reveal the role of neuronal *Mecp2* expression in the regulation of weight and metabolism in mouse models.

Interestingly, in contrast to the male *Mecp2^-/y^* mice, *Mecp2e1^-/+^* female mice did not show clear changes in either energy expenditure or food intake prior to increases in body weight in the C57BL6/J background. These results suggest that there is a gene dosage effect on metabolic phenotypes in this RTT model that is observed prior to and somewhat independent from body weight differences. More subtle, but nonsignificant differences in food intake or energy expenditure may be responsible for the weight gain in the female mutant mice, such as the increase in triglycerides observed in *Mecp2e1^-/+^* liver. More comprehensive studies of additional animals and time points are likely needed to determine the causes of weight gain in *Mecp2e1^-/+^* mice.

In conclusion, our results demonstrate that MeCP2-e1 deficiency is sufficient to cause hind clasping, motor, and metabolic phenotypes in both female and male mice. Increased body weight and fat mass was due to decreased energy expenditure, not increased food intake, in MeCP2-e1 deficient male mice. This metabolic phenotype is consistent across multiple RTT models of different genetic mutations and backgrounds and is associated with neuronal deficiency of MeCP2. Understanding how neuronal MeCP2e1 deficiency interacts with metabolism during the molecular pathogenesis of disease progression in female RTT models is therefore of critical importance to the successful design of future therapies.

## Materials and Methods

### Breeding and Cross Fostering

*Mecp2-e1* heterozygous females (*Mecp2e1^-/+^*) were maintained on a pure C57BL/6 J background by breeding to wild-type C57BL/6 J males (Jax strain 000664). To mitigate the impact of poor maternal care by *Mecp2e1^-/+^* dams, all pups were cross-fostered to CD1 foster dams within the first 48hours of birth. Briefly, *Mecp2e1^-/+^* females were paired with wild-type C57BL/6 J males for two weeks. CD1 male and female mice were also paired over the same time period so that litters would coincide. All males were then removed from the mating cages. Within the first 48 hours of birth the entire litter from *Mecp2e1^-/+^* dams were removed from their birth mother and placed with a CD1 dam that had given birth to a litter within the previous 24–48 hours. All but two the CD1 offspring were euthanized upon cross fostering and all pups remained with the CD1 dam until weaning at postnatal day (PND) 21. Mice were maintained in a conventional temperature controlled vivarium, with ad libitum access to food and water, with the lights on from 7 AM to 7 PM. Behavioral testing was conducted in adjacent testing rooms during the light phase of the circadian cycle. All experiments were conducted in accordance with the National Institutes of Health Guidelines for the Care and Use of Laboratory Animals. All procedures were approved by the Institutional Animal Care and Use Committee of the University of California, Davis and are covered under IACUC protocol #18881.

### Developmental milestones (PND 6–20)

Developmental milestones (75–77) were measured on PND 6, 8, 10, 12, 14, 16, 18, and 20 as previously described (16). All measures were conducted by an experimenter blind to genotype. Body weight and body length (nose to anus) were measured using a scale (grams) and ruler (cm). Pinnae detachment, ear detachment, eye opening, incisor eruption, and fur development were each rated on a scale of 0 (not present) to 3 (fully present). Cliff avoidance was tested by placing each pup near the edge of a clipboard, gently nudging it towards the edge, and scoring avoidance on rating scale from 0 (no edge avoidance) to 3 (complete turning and backing away from the edge). Righting reflex was tested by placing each pup on its back, releasing it, and measuring the time for it to fully flip over onto four paws for two trials on each developmental day. Failures to flip over in the righting test were recorded as a maximum score of 30 seconds. Grasping reflex was tested by brushing the forepaws with a cotton tipped applicator and rating the grasping reflex from 0 (none) to 3 (strong). Auditory startle was tested using a rating scale from 0 (no response) to 3 (large flinch and turn) in response to a finger snap near the pup’s head. Bar holding was tested by placing each pup’s front paws on a cotton tipped applicator stick and gently lifting up. Scoring consisted of a rating scale from 0 (immediate fall) to 3 (stay on and climb up). Level screen reflex was tested by placing each pup on a screen, gently pulling the tail, and rating the level of resistance (0 = none to 3= strong). Vertical screen reflex was tested by placing each pup on a screen at a 90 degree angle and rating the pup’s ability to remain on the screen from 0 (immediate fall) to 3 (hold on and climb to top). Negative geotaxis was tested by placing each pup, facing downwards, on a screen angled at 45 degrees from parallel, and scored on a scale from 0 (no turning) to 3 (complete turning and climbing to the top of the screen).

### Adult Behavioral Battery

Behaviors were assessed in young adult mice from PND 46 to152 in females and PND 35–95 in males. Mice had free access to food and water and lights were maintained on a 12:12-h light/dark cycle, with all behavioral testing performed during the light portion of the cycle. All behavioral assessments were conducted by an experimenter blinded to genotype.

### Body weight and Clasping Score (weekly PND 28–152 for females, PND 28–128 for males)

Weekly body weights and hind limb clasping were assessed beginning from PND 28. Hind limb clasping was rated on a scale from 0 (no clasping behavior) to 5 (both hind limbs tucked in and abdomen contracted)(16, 39).

### Elevated Plus-maze (PND 46–47 for females, PND 35 for males)

Each mouse was individually placed in the center area of a black Plexiglas automated elevated plus-maze (Med-Associates, St. Albans City, VT), under 300 lux illumination for a five minute test session (76, 78, 79). The entries per arm, total entries, and the time spent in each arm were recorded and compared between conditions (77).

### Light↔Dark Exploration (PND 49–50 for females, PND36 for males)

The light↔dark exploration task was performed as previously described (76). Briefly, the automated photocell-equipped apparatus was made of two Plexiglas compartments separated by a partition with a small opening. One compartment was transparent and illuminated by an overhead light (400 lux). The other compartment was made of black Plexiglas and closed on top. Each mouse was placed into the center of the light compartment and allowed to freely explore for 10 minutes. The number of transitions between light and dark sides and time spent in each compartment were recorded and compared between conditions.

### Open Field Exploration (PND 53–55 for females)

Exploratory behavior was assessed in a novel open field as previously described (76). Each mouse was placed in an automated VersaMax Animal Activity Monitoring System (AccuScan Instruments, Columbus, OH) and allowed to freely explore for 30 minutes. The number of horizontal and vertical beam breaks was used as a measure of horizontal and vertical activity respectively. Total distance traveled was used as a measure of total activity. Time spent in the center of the chamber compared to along the edges was recorded and compared between conditions.

### Gait Analysis (PND 60–61 and 144 for females, PND65 for males)

On the day prior to analysis all animals were habituated to restraint and paint application. On test day each animal was restrained, and the front feet were painted with blue while the back feet were painted with red paint (Sargent Art 22–3350 Washable Paint). The mouse was then allowed to walk down a straight alleyway lined with drawing paper (Strathmore). Colored footprints were analyzed for stride length (distance between successive forelimb and successive hind limb prints), hind-base (distance between the right and left hind prints), front-base (distance between right and left front prints), and paw separation (distance between the forepaw and hind paw placement)(80). Two to three footprints were analyzed per animal.

### Accelerating Rotarod (PND 62–64 and 140–143 for females)

Animals were given three trials per day with 1 hr inter-trial intervals, repeated over three consecutive days (76, 77). Each trial consisted of placing the animal on the rotarod apparatus (Ugo Basile mouse accelerating rotarod) and starting the rotation at 4 revolutions per minute (rpm). After the mouse was successfully walking, the rotarod was accelerated from 4 to 40 rpm over five minutes(76). The latency to fall from the rotating beam to the flange on the floor of the apparatus was recorded for each animal. The trial was terminated in cases where the mouse clutched the beam without walking, for five consecutive turns, or a maximum of 300 seconds had elapsed (16, 76, 77).

### Beam and Rod Walking (PND 88–92 and 152 for females, 60–64 and 95 for males)

A beam walking motor task was conducted as previously described (41). 59 cm long square dowels were suspended 68 cm above a cushioned landing pad. A goal box at the end of the beam consisted of a 12cm diameter cylinder to provide motivation to cross the beam. Each mouse was placed at the end of the beam and the time to cross to the goal box on the other end was measured. On the day prior to testing all animals were given four practice trials on the largest diameter square beam in order to become accustom to the procedure(80). On the test day each animal was sequentially tested on three square beams (24 mm, 19 mm, and 15 mm) and three round rods (17 mm,12 mm, and 9 mm). Testing sequence was based on presentations of decreasing diameter to present increasing levels of difficulty. Each mouse was given two trials on each beam, separated by approximately 30 min. The time to transverse the beam was recorded and averaged across the two trials for each beam. A maximum time of 60 sec was assigned to individuals that failed to cross the beam in that duration. In the small number of cases where mice fell from the beam, a score of 60 sec was assigned.

### Social Approach (PND 74–78 in females)

Sociability was assessed using the three chambered social approach task as previously described (77, 81, 82). The testing apparatus consisted of three connected chambers made of white matte Plexiglas, separated by two sliding doors. Prior to testing, the subject mouse was placed in the empty center chamber with the doors closed for 5 min. Doors were then removed to allow free access to all three empty chambers for 10 min. Distance traveled and the number of entries into the two side chambers were automatically scored using threepoint tracking with EthoVison XT tracking software (Noldus Information Technology Inc. Leesburg, VA). The mouse was then confined to the center chamber and a novel female 129Sv/ImJ mouse was placed in an inverted wire pencil cup in one chamber. An empty inverted wire pencil cup was simultaneously placed in the other chamber, to serve as the novel object. The partition doors were opened and the subject mouse was given free access to all three chambers for 10 min. Time spent and number of entries into each chamber, time spent sniffing the novel mouse and time the novel object, and the total distance traveled were simultaneously scored using EithoVision XT (77, 81, 83).

### Short-Term Memory Novel Object Recognition and Object Location Recognition (PND 66–71 and 81–85 respectively in females) and (PND 39–44 and 47–52 respectively in males)

Novel object recognition testing was performed as described previously (16, 40, 84). Briefly, all animals were handled for 2 min per day by the experimenter for three days. All animals were then habituated to the empty training arena for 10 min per day for four days. Distance travelled was recorded for each day of habituation and during training and testing. For the training session each animal was placed into the arena with two identical objects for 10 min. A 5 min testing session occurred 60 min after training. For Novel Object Recognition testing, one of the training objects was replaced with a novel object. For Object Location Recognition testing, one of the training objects was placed in a novel location. Time spent exploring each object was scored by an experimenter blind to all conditions. Animals exploring less than 3 seconds total for both objects during training or testing were excluded from further analysis (85).

### Energy expenditure, food intake and physical activity

On day 0 mice were transferred to single chamber housing in Comprehensive Laboratory Animal Monitoring System (CLAMS) chambers (Columbus Instruments, Columbus, OH) for acclimation. On day 1 mice in CLAMS chambers were transferred to a temperature controlled cabinet set to 24°C at 9:00 AM. Data recording was initiated with the following settings; initial reference reading (2 min) +2 min/cage, reference settle time:90 seconds, reference measure time:30 seconds, cage settle time:90 seconds, cage measure time:30 seconds, time to cycle for 8 cages:18 min (3 readings/hr). Data including volume of oxygen consumed (VO_2_), volume of carbon dioxide produced (VCO_2_), mass of food consumed, Total Number of X-axis IR-Beam Breaks (X-Tot), Number of Ambulatory (beam breaks in sequence) X-axis IR-Beam Breaks (X-amb), Total Number of Y-axis IR-Beam Breaks (Y-Tot), Number of Ambulatory Y-axis IR-Beam Breaks (Y-amb) and Number of Vertical (Rearing) Motions (Z-Tot) was acquired continuously for four days. The mice were acclimated to the CLAMS for 48 hours, and then data was analyzed from the subsequent 48 hours of CLAMS measurements. Respiratory Exchange Ratio (RER) was calculated as the ratio of carbon dioxide production/oxygen consumption. Heat production or energy expenditure is calculated from the equation, (Heat production = CV * VO_2_) where CV = 3.815 + 1.232 * RER. Calorimeter analyzer calibration was completed in the morning prior to each 24 hour measurement. A 0.50% CO_2_ and 20.50% O_2_ (balance nitrogen) calibration gas (Airgas, Sacramento, CA) and dry room air were used to calibrate the analyzers. At the start and end of the experiments, the performance of the entire calorimetry system was validated by bleeding a 20% CO_2_ and 80% N_2_ standard (Airgas, Sacramento, CA) into each calorimetry chamber at a regulated rate using an OxyVal gas infusion system (Columbus Instruments, Columbus, OH) and measuring recovery of CO_2_ and dilution of O_2_ in the chamber outflow.

### DEXA analysis

On day 5 mice were removed from CLAMS chambers at 9AM and euthanized by isoflurane inhalation prior to body composition analysis by Dual-energy X-ray absorptiometry (DEXA). Percent fat, total fat mass, bone mineral content (BMC), bone mineral density (BMD), bone area and subject length were calculated from absorption of the two X-ray frequencies. Mice were then necropsied immediately after DEXA with removal of brain, liver and spleen. One half of each tissue was flash frozen and stored at -80 degrees Celsius.

### Untargeted complex lipid analysis of liver

The untargeted lipidomics procedure began with lipid extraction from frozen, RNA later (Qiagen) preserved liver using methyl *tert*-butyl ether (MTBE) with the addition of internal standards, followed by ultra-high pressure liquid chromatography (UHPLC) on a Waters CSH column, interfaced to a Q-TOF mass spectrometer (high resolution, accurate mass), with a 15 min total run time. Data are collected in both positive and negative ion mode, and analyzed using Mass-Hunter (Agilent). Extraction of hepatic lipids was based on the “Matyash” method 36 (Matyash V, J. Lipid Research, 2008) which was subsequently modified. Briefly extraction is carried out using a bi-phasic solvent system of cold methanol, MTBE, and water. In more detail, cold methanol (225 μL) containing a mixture of odd chain and deuterated lipid internal standards [lysoPE(17:1), lysoPC(17:0), PC(12:0/13:0), PE(17:0/17:0), PG(17:0/17:0), sphingosine (d17:1), d_7_-cholesterol, SM(17:0), C17 ceramide, d_3_-palmitic acid, MG(17:0/0:0/0:0), DG(18:1/2:0/0:0), DG(12:0/12:0/0:0), and d_5_-TG(17:0/17:1/17:0)] is added to a 20 μL sample aliquot, which is placed into a 1.5 mL Eppendorf tube, and the tube is vortexed for 10 s. Then, 750 μL of cold MTBE containing CE (22:1) (internal standard) are added, followed by vortexing for 10 s. and shaking for 6 min. at 4ºC. Phase separation is induced by adding 188 μL of mass spec-grade water. After vortexing for 20 s. the sample was centrifuged at 14,000 rpm for 2 min. The upper organic phase was collected in two 300 μL aliquots. One was stored at – 20ºC as a backup and the other was evaporated to dryness in a SpeedVac. Dried extracts were resuspended using a mixture of methanol/toluene (9:1, v/v) (60 μL) containing an internal standard [12-[[(cyclohexylamino) carbonyl]amino]-dodecanoic acid (CUDA)] used as a quality control.

### LC/MS parameters

The LC/QTOFMS analyses were performed using an Agilent 1290 Infinity LC system (G4220A binary pump, G4226A autosampler, and G1316 C Column Thermostat) coupled to either an Agilent 6530 (positive ion mode) or an Agilent 6550 mass spectrometer equipped with an ion funnel (iFunnel) (negative ion mode). Lipids were separated on an Acquity UPLC CSH C18 column (100 × 2.1 mm; 1.7 μm) maintained at 65°C at a flow-rate of 0.6 mL/min. Solvent pre-heating (Agilent G1316) was used. The mobile phases consisted of 60:40 acetonitrile:water with 10 mM ammonium formate and 0.1% formic acid (A) and 90:10 propan-2-ol:acetonitrile with 10 mM ammonium formate and 0.1% formic acid. The gradient was established as follows: 0 min 85% (A); 0–2 min 70% (A); 2–2.5 min 52% (A); 2.5–11 min 18% (A); 11–11.5 min 1% (A); 11.5–12 min 1% (A); 12–12.1 min 85% (A); 12.1–15 min 85% (A). A sample volume of 3 μL was used for the injection. Sample temperature was maintained at 4°C in the auto-sampler.

The quadrupole/time-of-flight (QTOF) mass spectrometers were operated with electrospray ionization (ESI) performing a full scan in the mass range *m*/*z* 65–1700 in positive (Agilent 6530, equipped with a JetStreamSource) and negative (Agilent 6550, equipped with a dual JetStream Source) modes producing both unique and complementary spectra. Instrument paramaters are as follows (positive mode) Gas Temp 325°C, Gas Flow 8 l/min, Nebulizer 35 psig, Sheath Gas 350°C, Sheath Gas Flow 11, Capillary Voltage 3500 V, Nozzle Voltage 1000 V, Fragmentor 120 V, Skimmer 65 V. Data (both profile and centroid) are collected at a rate of 2 scans per second. In negative ion mode, Gas Temp 200°C, Gas Flow 14 l/min, Fragmentor 175 V, with the other parameters identical to positive ion mode. For the 6530 QTOF, a reference solution generating ions of 121.050 and 922.007 *m/z* in positive mode and 119.036 and 966.0007 m/z in negative mode, and these are used for continuous mass correction. For the 6550, the reference solution is introduced via a dual spray ESI, with the same ions and continuous mass correction. Samples were injected (1.7 in positive mode and 119.036 and 966.0007 m/z in nwith a needle wash for 20 seconds (wash solvent is isopropanol). The valve was switched back and forth during the run for washing; this has been shown to be essential for reducing carryover of less polar lipids.

### Untargeted Lipidomics Data Analysis

For data processing the MassHunter software was used, and a unique ID was given to each lipid based on its retention time and exact mass (RT_mz). This allows the report of peak areas/heights or concentration of lipids based on the use of particular internal standards. Lipids are identified based on their unique MS/MS fragmentation patterns using in-house software, Lipidblast. Using complex lipid class-specific internal standards this approach is used to quantify >400 lipid species including: mono-, di- and triacylglycerols, glycerophospholipids, sphingolipids, cholesterol esters, ceramides, and fatty acids. Peak area/heights (intensity) data was recorded from the MS/MS data.

### Metaboanalyst analysis

MetaboAnalyst 4.0 was used for analysis of QTOF-MS/MS peak intensity data. Settings for MetaboAnalyst (http://www.metaboanalyst.ca/) analysis were as follows; Data filtering using interquantile range, sample normalization by sum, data transformation using log, Pareto scaling. Sample groups were considered to be unpaired. For fold change analysis Mecp2-e1-/+ (MUTANT) peak heights over Mecp2-e1+/+ (WT) peak heights with a cutoff of log2 fold change (FC) at > 1.0 or < 1.0 was used.

### Statistical analyses of the data

Male and Female developmental milestone data were analyzed separately. Continuous outcomes (body weight, body length, average righting reflex time) were analyzed using linear mixed effects models, including fixed effects for genotype, time, and litter size (number of pups), the interaction genotype x time interaction, and random intercepts for mouse and litter (nlme and lsmeans R packages). Categorical ratings of pinnae detachment, eye opening, incisor eruption, and fur development were analyzed using mixed effects Cox proportional hazards models of the time to full development; these models included fixed effects for genotype and litter size and a random slope for litter (coxme and car R packages). Categorical ratings of cliff avoidance, auditory startle, negative geotaxis, bar holding, vertical screen, and level screen were analyzed using global F tests from regression models for ordinal data via cumulative link (mixed) models (clmm function from Ordinal R package) with fixed effects for genotype, time, and litter size as well as the genotype x time interaction and random intercepts for mouse and litter. For males auditory startle ratings on the first time of assessment were not variable enough for inclusion in the model. For level screen scores, there was insufficient variability in scores for males beyond day 14 and for females beyond day 12 so only the first four or three time points were analyzed respectively.

Post-weaning weight was analyzed separately for each sex using a linear mixed effects model, including fixed effects for genotype, time, genotype x time interaction and random effect of mouse. Benjamini-Hochberg corrected post hoc comparisons were made for each time point. The development of significant hind limb clasping in the mutant mice was assessed for each timepoint where at least one animal showed a non-zero clasping score. At each timepoint examined the number of animals with non-zero clasping scores was assessed using a binomial test (R package binom, with the binom.confint function with two.sided test) to assess if clasping was significantly greater than zero (no clasping) with an overall Benjamini-Hochberg correction across time points.

All other behavioral analyses included parametric independent, two tailed t-tests or ANOVA models with Benjamini-Hochberg corrected multiple posthoc tests. Repeated measures tasks were accessed using a mixed model repeated measures ANOVA with a repeated measure and genotype and fostering as factors. All analyses were conducted using R, version 3.3.1(86) or prism version 7. Linear mixed effects modeling was conducted using the R package lme4, version 1.1–12(87), or nlme, version 3.1–131.1(88) with lsmeans, version 2.27–61 (89), mixed effects Cox proportional hazards modeling was conducted using the R package coxme, version 2.2–5(90), and mixed effects proportional odds logistic regression modeling was conducted using the R packages ordinal, version 2015.6–28 (91).

For CLAMS and DEXA analysis all data are expressed as mean ± SEM unless otherwise indicated. Differences within a group were evaluated by t-test while differences between genotypes and sexes were evaluated using a two-way analysis of variance (92). Post hoc analysis was carried out using a Tukey’s honest significant difference test to determine which diet and age groups differed significantly. For energy expenditure data, analysis of covariance was used with either body weight or lean mass as a covariate in the model.

## Acknowledgements

AVC, DHY, JML and JNC designed experiments and wrote the manuscript. AVC and MP conducted behavioral experiments. BDJ conducted statistical analyzes. AVC, AC, AN, and DHY assisted with behavioral scoring and data analysis. JLR and JR performed the CLAMS and DEXA analysis. Metabolomics analysis was conducted by West Coast Metabolomics Center. The EE ANCOVA analysis was provided by the NIDDK Mouse Metabolic Phenotyping Centers (MMPC, www.mmpc.org) using their Energy Expenditure Analysis page (http://www.mmpc.org/shared/regression.aspx). Thank you to Heather Boyle and the animal care staff at UC Davis for colony management, breeding and cross fostering.

